# The anger-insula-body link in heart arrhythmias

**DOI:** 10.64898/2026.10.19.760035

**Authors:** Viridiana Mazzola, Elisabeth Martins-Dominati, Guido Bondolfi, Giampiero Arciero

## Abstract

Anger is a well-known risk factor in cardiovascular diseases (CVDs) since it triggers a sympathetic hyperactivity of peripheral autonomic nervous system (ANS). As our previous findings showed, facing an angry situation instead of a joyful one elicits in healthy participants an affective engagement that involved mainly the insular reactivity and its connectivity. On the other hand, stimulating the insula, the key neural hub of the cardiovascular brain, can induce cardiac arrhythmias. Unfortunately, still little is known about the interplay between the anger and the insula as the underlying emotional-brain-body link that can trigger or precipitate a series of neurogenic cardiac events in CVDs patients. To address this question, we employed a multimodal method combining neuroimaging, behavioral and blood data to gain a more complete picture of the insula activity and its connectivity in patients with different types of heart arrhythmias. The present findings show a greater generalized involvement of the bilateral posterior insulae in the patients group both when facing angry context and squeezing the handgrip device, suggesting an automatic engagement of the sympathetic nervous system/SNS. Moreover, the bilateral insulae were functionally coupled with the bilateral lateral ventricles selectively under angry conditions, suggesting an insular role in glymphatic dysfunction and CSF dynamics due to the neurovascular coupling. Clearly, understanding this anger-insula-heart link may factually contribute to lower the probability of adverse cardiac outcomes in orienting the necessary treatment of the person, taking into account the emotion-related cardiovascular pathologies and the psychological health.

## Introduction

Anger is a well-known risk factor in cardiovascular diseases (CVDs). Indeed, it triggers a sympathetic hyperactivity of peripheral autonomic nervous system (ANS), therefore increasing CVDs in initially healthy populations but also worsening the prognosis in the patients with existing CVDs (Chida et al. 2009, Levine et al. 2021). In addition, anger facilitates those high-risk behaviors such as poor diet, less physical activity, smoking, poor sleep, or lower treatment adherence, which take unnoticeably root in everyday life. Consequently, the medical monitoring, prevention and cardiac rehabilitation programs aim to limit, change and re-educate these behaviors. But such an approach to mediate these high-risk behaviors via a behavioral pathways does not target this sympathetic hyperactivity as the underlying risk factor. Despite the abundance of medical knowledge in this field, CVDs are still a leading contributor to morbidity and mortality globally.

Actually, how anger experience may be linked to the cardiovascular brain circuit and the downstream physiological changes is not fully understood and the precise nature of these observed relationships has not yet been fully established.

The cardiovascular brain circuit is a complex multidimensional network and plays a central role in orchestrating the neurovascular, the neuroimmune, the central and peripheral autonomous systems in response to interoceptive and exteroceptive stimuli (Mohanta et al 2023). As a key neural hub of this circuit, the insula directly modulates ANS activity by influencing both sympathetic and parasympathetic branches of the brain-heart axis, thus regulating the neurocardiovascular balance in its being emotionally involved in our daily lives and in clinical conditions too. Stimulating the insula can induce cardiac arrhythmias as well as structural damage to the cells of the heart (myocytolysis) (Oppenheier et al 1991). It has even been suggested that the insula may be involved in acute emotion-induced cardiac alterations, arrhythmias, and sudden death, including Takotsubo cardiomyopathy (Ueyama, 2004). As our previous findings showed, facing an angry situation instead of a joyful one elicits in healthy participants an affective engagement that involved mainly the insular reactivity and its connectivity (Mazzola et al. 2016, 2020). However, this evidence relies on studies on individuals without CVDs. Unfortunately, still little is known about the interplay between anger and the insula as the underlying emotional-brain-body link that can trigger or precipitate a series of neurogenic cardiac events in CVDs patients.

To address this question, we employed a multimodal method combining different data and analyses to gain a more complete picture of the insula activity in CVDs patients. We employed our previous functional magnetic resonance imaging (fMRI) task on a sample of twenty patients with different types of heart arrhythmias and twenty healthy participants (Mazzola et al. 2016, 2020). All the participants watched angry and joyful situations during the fMRI scanning session. They watched two runs of video clips of an actor and an actress performing a daily act like grasping an object with different emotional expressions (neutral, joyful, or angry). In order to set two distinct emotional contexts, one run included only neutral and joyful expressions, while the other featured only neutral and angry expressions. We hypothesized that angry situations would show a higher involvement of the insula in patients compared to healthy controls. Then, to better understand the insula engagement during the angry situation, we used a psycho-physiological interaction (PPI) analysis to examine changes in the functional connectivity of the insular region under the angry run (angry vs. joyful run). In addition, since the isometric handgrip can also be used to investigate the underlying mechanisms of sympathetic nervous activity, we checked for any significant group differences while performing a dynamic handgripping task during the fMRI session. To check for any significant differences in the baseline insular activity between groups, we collected both BOLD and arterial spin labeling (ASL) data in resting state.

## Methods

### Ethics Statement

The present study was approved by the local ethical committee. Informed written consent was obtained from all participants before the inclusion in the study. The authors assert that all procedures contributing to this work comply with the ethical standards of the relevant national and institutional committees on human experimentation and with the Helsinki Declaration of 1975, as revised in 2008.

### Participants

Twenty patients who received a diagnosis or were treated for different types or arrhythmia with or without catheter ablation in the last 3 months at the Cardiac Pacing Unit, Cardiology Department, University Hospitals of Geneva (HUG) (Table 1), were recruited from 1/03/2017 to 15/12/2019 by our nurse (coauthor E. Martins-Dominati) as well as twenty age- and gender-matched healthy controls (patients group: 11 females: median age 39 years old, range 17–62; 9 males: median age 49 years old, range: 18-64; health group: 11 females, mean age 41 years old, range 17–65; 9 males median age 48 years old, range: 19–63). Exclusion criteria included a history of drug or alcohol abuse, previous head trauma with loss of consciousness, pregnancy, and substantial medical comorbidities or significant psychiatric conditions. All the participants had more than 16 years of schooling. After a medical history interview conducted by our nurse, participants were asked to describe the way they usually get angry, namely, what is the trigger (situations), the level of involvement of bodily states in terms of higher-medium-low intensity, how long their anger lasts and how it ends (duration and regulation) (Mazzola et al. 2020). Then, following a semi-structured interview about stressful life events, they were asked to assess stressors in the form of ‘events’ (discrete episodes) and ‘difficulties’ (lasting at least 2 weeks) during the year before symptom onset. For healthy controls, the 2 years prior to the interview date were studied to minimize differences in time from interview to life events, and therefore reliability of recall, compared to patients.

**Table 1:**
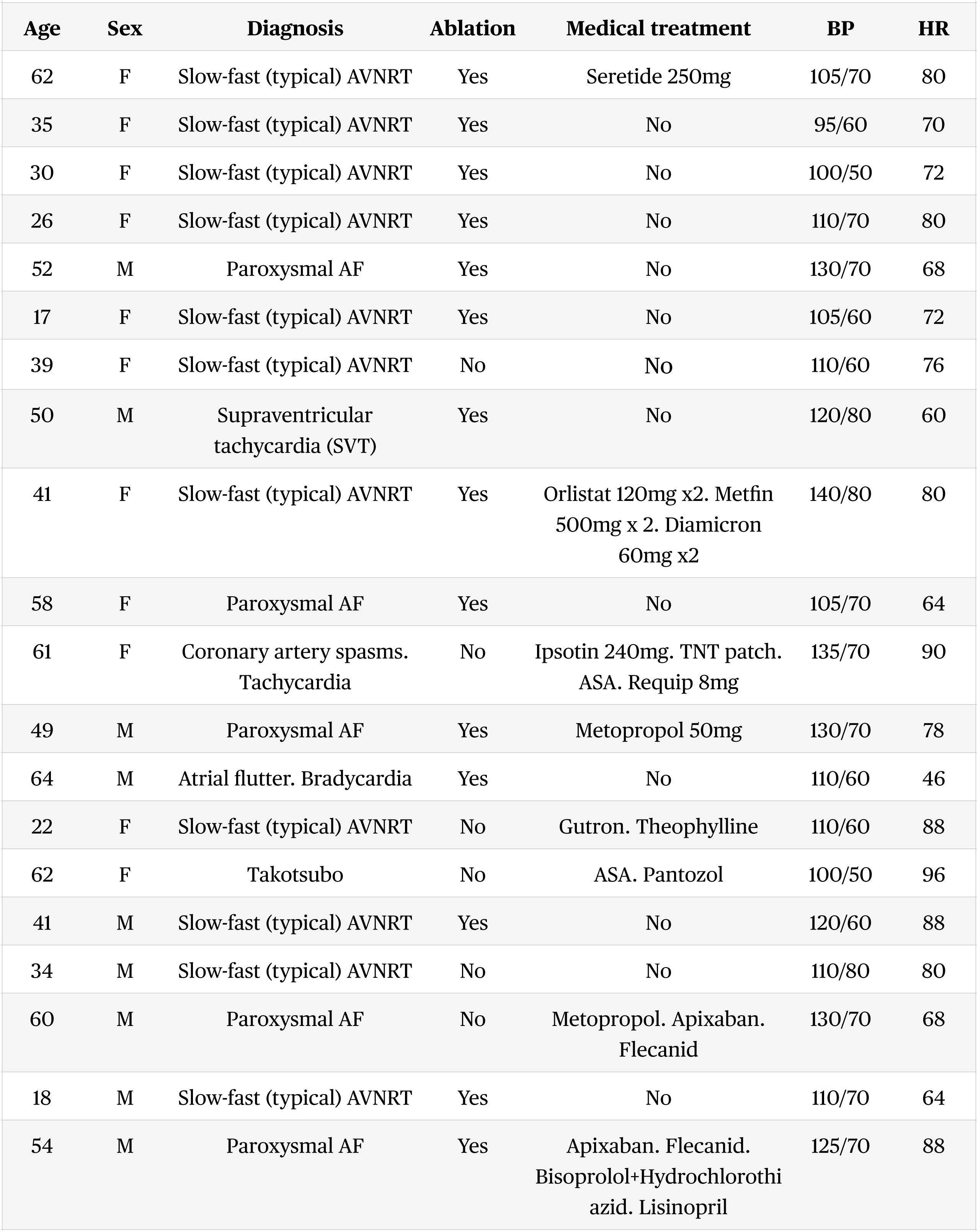
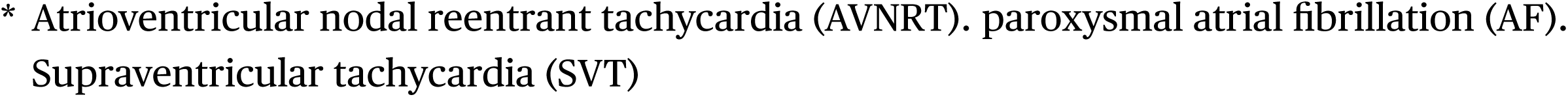
Patients information.

| Age | Sex | Diagnosis | Ablation | Medical treatment | BP | HR |
| --- | --- | --- | --- | --- | --- | --- |
| 62 | F | Slow-fast (typical) AVNRT | Yes | Seretide 250mg | 105/70 | 80 |
| 35 | F | Slow-fast (typical) AVNRT | Yes | No | 95/60 | 70 |
| 30 | F | Slow-fast (typical) AVNRT | Yes | No | 100/50 | 72 |
| 26 | F | Slow-fast (typical) AVNRT | Yes | No | 110/70 | 80 |
| 52 | M | Paroxysmal AF | Yes | No | 130/70 | 68 |
| 17 | F | Slow-fast (typical) AVNRT | Yes | No | 105/60 | 72 |
| 39 | F | Slow-fast (typical) AVNRT | No | No | 110/60 | 76 |
| 50 | M | Supraventricular tachycardia (SVT) | Yes | No | 120/80 | 60 |
| 41 | F | Slow-fast (typical) AVNRT | Yes | Orlistat 120mg x2. Metfin 500mg x 2. Diamicron 60mg x2 | 140/80 | 80 |
| 58 | F | Paroxysmal AF | Yes | No | 105/70 | 64 |
| 61 | F | Coronary artery spasms. Tachycardia | No | Ipsotin 240mg. TNT patch. ASA. Requip 8mg | 135/70 | 90 |
| 49 | M | Paroxysmal AF | Yes | Metopropol 50mg | 130/70 | 78 |
| 64 | M | Atrial flutter. Bradycardia | Yes | No | 110/60 | 46 |
| 22 | F | Slow-fast (typical) AVNRT | No | Gutron. Theophylline | 110/60 | 88 |
| 62 | F | Takotsubo | No | ASA. Pantozol | 100/50 | 96 |
| 41 | M | Slow-fast (typical) AVNRT | Yes | No | 120/60 | 88 |
| 34 | M | Slow-fast (typical) AVNRT | No | No | 110/80 | 80 |
| 60 | M | Paroxysmal AF | No | Metopropol. Apixaban. Flecainid | 130/70 | 68 |
| 18 | M | Slow-fast (typical) AVNRT | Yes | No | 110/70 | 64 |
| 54 | M | Paroxysmal AF | Yes | Apixaban. Flecainid. Bisoprolol+Hydrochlorothiazid. Lisinopril | 125/70 | 88 |

\* Atrioventricular nodal reentrant tachycardia (AVNRT). paroxysmal atrial fibrillation (AF). Supraventricular tachycardia (SVT)

All the participants completed the State-Trait Anger Expression Inventory for Adult (STAXI-2) (Spielberger 1999) in order to control for their general tendency to experience anger, the Body Perception Questionnaire (BPQ) (Porges, 1983), the State-Trait Anxiety Inventory (STAI-ST), and Beck Depression Inventory (BDI-II).

Each participant was visited us three times over two months. During the first visit, the participant will have the clinical psychological interview, the medical history interview, and the administration of the questionnaires. The modes of data collection by questionnaires were: self-administered in an electronic format (computer mouse/keyboard) and computer assisted face-to-face interviewing for the medical history. During the second visit, the participant underwent BP measurement, blood sampling (Hémogard vert 4ml and Hémogard jaune 5 ml), and fMRI session. The blood sample was stored in the ice and transported to the laboratory of Toxicology at the HUG. For each participants, except one healthy participant, the plasma level of adrenaline, noradrenaline, dopamine, and serotonine was obtained. Due to technical problems, only the blood samples of 12 patients and 16 healthy participants were analyzed by the V-Plex Proinflammatory Panel 1 human kit from the MesoScale Discovery (MSD) platform (Rockville, MD, USA).

### Self-report questionnaires

Questionnaire data were analyzed using JASP (2025). Mann-Whitney U tests between groups were performed to test for the group differences effect on STAXI-2, Body Perception Questionnaire (BPQ), BDI and STAI-T and S. We reported the exact significance level of p<0.001 (two-tailed). In order to test for any significant correlation between STAXI and BPQ, we performed Spearman’s rho correlation analyses. We reported the exact significance level of p<0.001 (one/two-tailed).

### fMRI task

The fMRI task employed was described in detail in previous studies (Mazzola et al., 2013, 2016, 2020). Briefly, the functional MRI session consisted of two successive scanning runs with an event-related design. Each run included one emotion ( joy or anger), plus the neutral facial expression, and consisted of four experimental view conditions. All visual stimuli consisted of video clips of 1.3 seconds. There were 4 different video conditions in each run, showing the following actions: the trunk with an arm grasping an object on a table (acting alone), a person with a neutral facial expression grasping an object on a table (neutral acting), a person with a joyful or angry facial expression grasping an object on a table ( joyful acting in the joyful run or angry acting in the angry run), and a joyful or angry dynamic facial expression without any acting action ( joyful face in the joyful run or angry face or in the angry run). The recording and editing of videos were made using the Blue Screen technique in order to superimpose on the same trunk different emotional facial expressions. Thereby, we investigated the brain activity elicited by the observation of someone acting in angry and joyful situations, while keeping action kinematics constant. Two professional actors, a female and a male, were enrolled as models for the videos. The everyday objects to be grasped were put on a table, e.g., phone, pen, keys, bottle, cup, and glass. To circumvent any motor interference, we used a passive viewing task, and participants were instructed to remain still without performing any movement, and to avoid any imitation or mental imagery of the actions shown, and to carefully look at the video clips in order to get involved. The presentation order of the two runs was counterbalanced across subjects. In each scanning run, there were 128 visual stimuli presented in random order, with an average 1810 ms interstimulus interval (ISI). Thirty-eight additional null events, each lasting 2700 ms, contributed to randomly jitter the stimulus onsets. Total scanning run time was about 16 minutes.

The handgrip task consisted of about 4 min of dynamic handgripping, i.e., repeated 15 s right hand contractions alternated with 15 s of relaxation. Participants were instructed to squeeze the handgrip with their right hand to their maximum ability when seeing the sign “on” on the screen and to relax when the sign “off” appeared.

### fMRI scan

All the subjects underwent fMRI session between the hours of 10:00 and 13:00 to control for the effects of circadian rhythm at the Brain and Behavior Laboratory (BBL) at the Medical Center of Geneva (CMU) using a 3.0 Siemens Magnetom TrioTim scanner equipped with a 32-channel head coil. All functional data were acquired using a gradient-echo planar imaging (EPI) sequence using T2* weighted blood oxygenation level dependent (BOLD) contrast, with 35 interleaved axial slices (3 mm thick) encompassing the entire cerebrum and the cerebellum (TR 2 s; TE 30ms; FOV 24 cm; a isotropic voxel size of 3 × 3 × 3 mm). A total of 150 EPI volume images were acquired for the resting state, a total of 235 EPI volume images for each session of the angry-joyful task, whereas the hand grip task featured a total of 93 EPI volume images.

A high-resolution T1-weighted anatomical scan was performed for each participant (MPRAGE, TR=1900ms, TE=2.27ms, TI=900ms, flip angle=9°, 192 sagittal slices, voxel dimensions=1mm isotropic, 256×256 pixels, PAT factor=2).

Pseudo-continuous ASL (pcASL) images were acquired using an echo-planar imaging (EPI) sequence with the following parameters: repetition time (TR) 3,500 ms, echo time (TE) 10 ms, slice thickness 5 mm without gap, field of view (FOV) 224 mm × 224 mm, matrix size 128 × 128, inversion time (TI_1_ 700 ms, TI_2_ 1800 ms), number of excitations 1, number of slices 20, total scan time 5 min and 31 s. ASL imaging protocol acquired an M_0_ image and 45 control/label pairs. Bio-signal variables, including HRV, PPG and respiration rate, were recorded using a Biopac MP150 (Biopac module, Biopac Systems Inc., USA). The photoplethysmogram (PPG) was recorded via an optical sensor attached to the distal phalanx of the left index finger. Specifically, the electrocardiogram (ECG) signal was recorded using an ECG100C module with lead II to analyze HRV, and the impedance cardiography signal was recorded using a NICO100C module to calculate the stroke volume (SV) and CO. The PPG signal was acquired using a PPG100C module to determine the average amplitude and average systolic area, and the RSP signal was acquired using an SKT100C module to calculate the respiration rate. The preprocessing and frequency domain HRV analysis of PPG data were run using the automated routine of the Acqknowledge 4.2 software.

### fMRI data preprocessing

Data were preprocessed and analyzed using statistical parametric mapping SPM12 v7771 (Wellcome Department of Cognitive Neurology, London, UK), implemented in MatLab R2023b (MathWorks^TM^). A fixed-effect model at a single-subject level was performed to create images of parameter estimates, which were then entered into a second-level random-effects analysis. For each subject, functional images were first slice-timing corrected, using the middle slice acquired in time as a reference, and then spatially corrected for head movement, using a least-squares approach and six-parameter rigid body spatial transformations. High-resolution anatomical T1 images were coregistered with the realigned functional images to enable anatomical localization of the activations. The two runs were then entered as multiple sessions as implemented in SPM12. Structural and functional images were spatially normalized into a standardized anatomical framework using the default EPI template provided in SPM12, based on the averaged-brain of the Montreal Neurological Institute and approximating the normalized probabilistic spatial reference frame of Talairach and Tournoux [http://brainmap.org/icbm2tal/]. Functional images were spatially smoothed with a three-dimensional Gaussian filter (10mm full-width at half-maximum). The time series was temporally filtered to eliminate contamination from slow drift of signals (high-pass filter, 128 s) and corrected for autocorrelations using the AR(1) model in SPM12.

### General linear model

We performed two parallel but identical statistical analyses on the functional data for the whole-brain and cerebellar normalized images. Four event types were defined per subject per scanning run, corresponding to each condition of interest. In the joyful run, the conditions of interest were: acting alone, neutral acting, joyful acting, joyful face. In the angry run, the conditions were: acting alone, neutral acting, angry acting, angry face. Eight contrast images corresponding to these conditions from individual participants were entered at the second level into a repeated-measures 2 (groups) × 2 (sessions) × 4 (conditions) ANOVA (flexible factorial design implemented in SPM12). We reported regions that survived a threshold of P < 0.05 cluster-level FWE corrected, cluster size K = 8, as implemented in SPM12. Cerebral MNI coordinates were converted to the Talairach coordinate system by icbm2tal [http://brainmap.org/icbm2tal/]. Anatomic and Brodmann areas labeling of cerebral activated clusters was performed with the Talairach Daemon database [http://www.talairach.org/] and SPM Anatomy Toolbox [<u>41</u>]. We used the PhysIO Toolbox for preprocessing of physiological recordings and model-based noise correction of fMRI data using peripheral measures of respiration and cardiac pulsation. We used a separate normalization process for data from the cerebellum. The registration between individuals and MNI space is suboptimal in the cerebellum when using a standard whole-brain normalization process. Because cerebella vary relatively little between individuals compared with the cortical landmarks used for whole-brain normalization, it is possible to achieve a much better registration by normalizing the cerebella separately. Moreover, precise spatial registration is important because cerebellar structures are small compared to cortical structures. To this aim, we used the SUIT toolbox for SPM12, allowing us to normalize each individual’s structural scan to an infratentorial template, and then used the resulting deformation maps to normalize the cerebellar sections of each person’s functional images. The SUIT toolbox has the additional advantage that coordinates can be adjusted from MNI space to the corresponding coordinates on the unnormalized Colin-27 brain, which is described anatomically in a cerebellar atlas. We used this feature to identify anatomical regions within the cerebellum.

### Resting state fMRI

Functional images were preprocessed and denoised using the default options in CONN version 22a (Nieto-Castanon and Whitfield-Gabrieli, 2022). Arterial spin labeling (pcASL) images were processed using the Signal Processing Toolbox (ASLtbx) software (Zhu et al., 2018).

## Results

### Self-report questionnaires

The results about the STAXI-2 questionnaire showed a strong positive correlation in the patients group between the anger control-out and anger control-in scales (rho=0.766 p<0.001), whereas in healthy controls a very mild correlation between anger trait and anger in scale (rho=0.604, p<0.01).

The BPQ questionnaire revealed a strong positive correlation in the patients between the awareness and the autonomic nervous system/ANS scales (rho=0.751 p<0.001) in the patients, whereas in the healthy controls between the stress response/SR and ANS scales (rho=0.756 p<0.001), as well as between ANS and anger trait scales (rho=0.713 p<0.001).

As the interviews on anger management revealed, at the apex of their own anger, the majority of the patients reported to perceive a general involvement of their body (in terms of the urge to act also against the person or the situation, feeling very hot, increasing transpiration up to sweat, heart beating or changing the rhythm of the respiration, either faster or slower up to be “blocked”) that lasted for a time interval in between 30/45 minutes and some days according to the relevance of the episode. On the other hand, the majority of the healthy participants reported that they usually did not get angry but rather irritated, at their peak they did not report to perceive any significant change in bodily states, and the peak lasted from 5 to 15/20 minutes in the most relevant episodes.

About the semi-structured interviews on stressful life events, our patients reported to have undergone more stressful events in the last one-two years compared to healthy controls. Normative (not the common traumatic events) stressful events (i.e., events that happen to most of us sometime in our lifetime, such as a job loss or the end of an important relationship) meant for them events with an heavy and lasting emotional impact (i.e. feeling powerless, blocked, trapped, angry, alarmed), as well as a bodily impact (i.e. feeling more tired, worsened quality of sleep, muscle tension). The ratio of angry/alarmed to joyful events was more shifted towards the angry in the patients group.

### Blood test

No significant mean difference in catecholamines and serotonin blood level was found between groups, neither in CRP. Instead, a significant lower mean in the healthy controls than in the patients was detected in the IL12-IL23 (t=-2.343/U=25 p=0.015/0.008 T-test one tailed), the IL16 (U=36 p=0.048 T-test one tailed), and the PIGF (t=-2.162/U=25 p=0.021/0.008 T-test one tailed).

### Neuroimaging Results

#### Whole-brain fMRI results

Across sessions, the two groups showed significant differences. The patients group activated more the right posterior insula, the right putamen, the right amygdala, among other regions involved in the emotion-action processing as the precentral gyrus, the left inferior frontal gyrus/ IFG and the superior temporal gyrus according to our previous findings (P<0.05 FWE corrected; Table 2) (Fig.1). On the other hand, the healthy controls group showed a higher activation of the thalamus, the superior parietal lobule, the anterior cingulate cortex/ACC and the caudate, all regions related to the action processing (P<0.05 FWE corrected; Table 2) (Fig.1). Across groups, compared to the joyful one, the angry session activated more the right middle temporal gyrus and the right superior frontal gyrus, whereas the reverse contrast did not show anything significant at the same statistical threshold (P<0.05 FWE corrected; Table 2) (Fig.1).

**Fig. 1:**
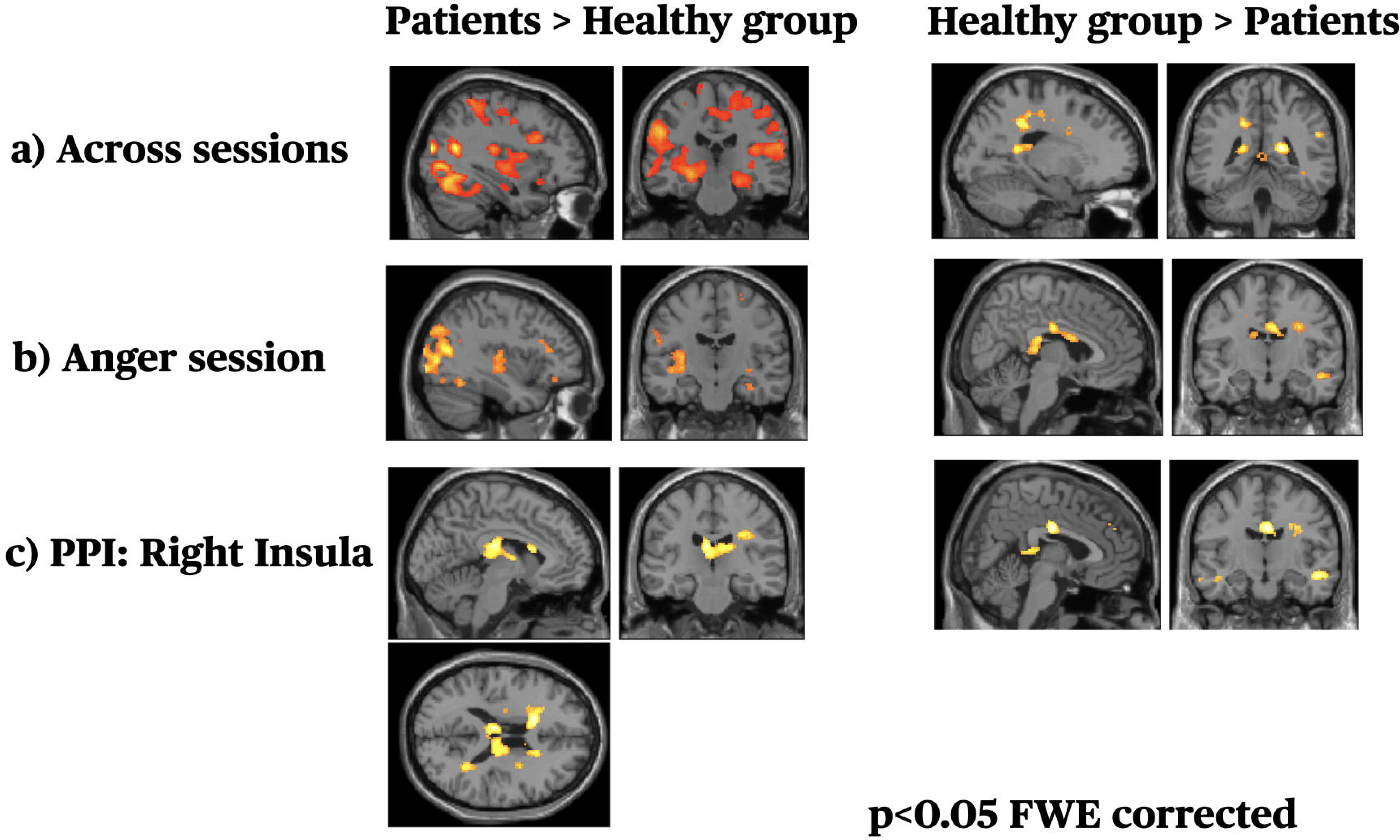
a) Brain sections (image threshold at p<0.05 FWE corrected) of the BOLD response of the main effects of group factor across sessions; b) brain sections (image threshold at p<0.05 FWE corrected) of the BOLD response of the main effects of anger factor between groups; c) brain sections of the BOLD response of the main effects of anger factor between groups in the PPI analysis with the right insulation as seed region (image threshold at p<0.05 FWE corrected).

**Table 2:** Anger/Joyful task. Whole-brain General Linear Model analyses. P<0.05 FWE.

|  | MNI |  |  |  |  |
| --- | --- | --- | --- | --- | --- |
| Region | x | y | z | Ke | Z Scores |
| HC group > PT group across sessions |  |  |  |  |  |
| R Precuneus | 18 | -42 | 16 | 1384 | 70.7 |
| L MCC | -18 | -42 | 38 | 431 | 5.90 |
| R MCC | 6 | -12 | 30 | 350 | 5.52 |
| R Precentral Gyrus | 30 | -12 | 32 | 175 | 5.36 |
| R Inferior Parietal Lobule Area PGa (IPL) | 42 | -56 | 48 | 15 | 5.34 |
| PT group > HC group across sessions |  |  |  |  |  |
| L Middle Temporal Gyrus | -54 | -66 | 16 | 10246 | Inf |
| R Precentral Gyrus | 54 | -2 | 44 | 191 | Inf |
| R Superior Temporal Gyrus Area TE 3 | 64 | -10 | 2 | 679 | Inf |
| L Precentral Gyrus | -48 | 8 | 36 | 395 | Inf |
| R Postcentral Gyrus Area 2 | 30 | -46 | 60 | 403 | Inf |
| L SMA | -30 | -10 | 52 | 681 | 7.45 |
| L Superior Parietal Lobule | -22 | -62 | 50 | 382 | 7.36 |
| L IFG (p. Orbitalis) | -24 | 30 | -12 | 64 | 7.18 |
| R IFG (p. Opercularis) | 46 | 18 | 28 | 222 | 7.13 |
| R Insula | 38 | -26 | 14 | 69 | 6.72 |
| L putamen | -22 | 12 | -2 | 51 | 6.25 |
| R Postcentral Gyrus Area 1 | 64 | -12 | 34 | 71 | 6.18 |
| R Hippocampus | 34 | -14 | -2 | 351 | 6.14 |
| L Temporal pole superior | -56 | 8 | -2 | 38 | 5.66 |
| R Precentral Gyrus | 24 | -18 | 66 | 45 | 5.36 |
| R Cerebelum (Crus 1) | 38 | -74 | -25 | 713 | Inf |
| R Cerebelum 6 | 24 | -66 | -27 |  | 7.27 |
| **Anger > Joyful run across groups** |  |  |  |  |  |
| R middle temporal gyrus | 22 | 26 | 46 | 148 | 3.82 |
| R superior frontal gyrus | 24 | 12 | 48 |  | 3.63 |
| **Joyful run HC group > PT group** |  |  |  |  |  |
| R Thalamus | 12 | -30 | 20 | 143 | 4.35 |
| **Joyful run PT group > HC group** |  |  |  |  |  |
| L Middle Temporal Gyrus | -54 | -66 | 16 | 28833 | Inf |
| L Middle Orbital Gyrus | -24 | 30 | -12 | 131 | 5.46 |
| R Thalamus | 12 | -16 | 4 | 74 | 4.82 |
| L putamen | -22 | 14 | -2 | 152 | 4.49 |
| R Middle Frontal Gyrus | 28 | 42 | 22 | 170 | 4.31 |
| R Cerebelum 6 | 30 | -72 | -23 | 107 | 4.89 |
| R Cerebelum (Crus 1) | 36 | -84 | -33 |  |  |
| **Angry run HC group > PT group** |  |  |  |  |  |
| R Mid Cingulate Gyrus | 4 | -12 | 30 | 188 | 5.90 |
| R Temporal Lobe | 48 | -16 | -16 | 252 | 5.48 |
| R Medial Frontal Gyrus | 18 | 26 | 32 | 154 | 5.29 |
| R Middle Temporal Gyrus | 48 | -48 | 0 | 135 | 4.69 |
| R IFG (p. Triangularis) | 48 | 22 | 12 | 113 | 4.51 |
| R SupraMarginal Gyrus Area PF (IPL) | 64 | -34 | 24 | 118 | 4.48 |
| L Postcentral Gyrus | -26 | -32 | 46 | 212 | 4.40 |
| **Angry run PT group > HC group** |  |  |  |  |  |
| L Middle Temporal Gyrus | -46 | -52 | 12 | 2779 | Inf |
| R Middle Temporal Gyrus | 56 | -62 | 12 | 3470 | 7.65 |
| R Precuneus | 28 | -76 | 40 | 441 | 6.42 |
| R Cerebelum (Crus 1) / Area FG2 | 46 | -60 | -20 | 271 | 5.58 |
| L Postcentral Gyrus Area PFt (IPL) | -54 | -26 | 32 | 414 | 5.29 |
| L Claustrum | -32 | -18 | 8 | 389 | 4.80 |
| L putamen | -32 | -16 | -2 |  | 4.45 |
| L Insula | -42 | -16 | -4 |  | 4.18 |
| R Cerebelum (Crus 1) | 38 | -74 | -25 | 37 | 5.31 |
| R Cerebelum 6 | 28 | -68 | -31 |  |  |

To address our research question, namely whether the processing of anger impacts more on the insula activity in the patients group, we tested the main effect of the angry session between groups. We found greater activation in the left posterior insula, as well as in the precuneus, the cerebellum crus I, and the putamen in the patients group. On the other hand, the healthy control group activated more the right middle cingulate cortex/MCC, the right IFG, and the supramarginal gyrus (P<0.05 FWE corrected; Table 2) (Fig.1). On the contrary, the insula did not reach a significant activation in the joyful session when we compared the two groups, indicating a specific role played in the angry session in the patients group (Table 2). PPG analyses did not show statistically significant differences between groups undergoing the angry-joyful task. Consistent with our a priori hypothesis, these results added support to the view that anger processing enhances activity within the insular region in the patients group.

We checked for the impact of the physiological noise correction on the previous findings. Interestingly, we did not find differences between groups in the bilateral posterior insulae activation across sessions and in the anger session at the same statistical threshold, but at a lower one (small volume correction/SVC p< 0.001), when the data were preprocessed with the physiological noise correction.

Next, we explored the changes in connectivity associated with these angry context effects focused on the right and left posterior insula, and RMCC. First, we used PPI to examine changes in the functional connectivity of the right posterior insula as a seed region under the angry context between groups. PPIs results showed that in the patients group this region was functionally coupled with the left lateral ventricle, the right anterior cingulate cortex/ACC, and the left caudate during both the angry session and the angry grasping (p< 0.05 FWE corrected; Table 3) (Fig.1). On the other hand, in the healthy group, the right posterior insula was functionally coupled with the left precentral gyrus, the right superior temporal gyrus, the left insula, and the right ACC, during both the angry session and the grasping angry (p< 0.05 FWE corrected; Table 3) (Fig.1). On the contrary, in the joyful context, no activation reached the statistical significance in both groups. The left posterior insula showed a functional coupling with the left insula and the right ventricle in the patients group under the angry context (p< 0.05 FWE corrected; Table 3), whereas in the healthy group it was more diffusely coupled, strongly with the bilateral MCC, the left SMA (p< 0.05 FWE corrected; Table 3). Under the joyful context, instead, the patients group showed a functional coupling with the right insulation, the right middle temporal and the left fusiform gyri, whereas the healthy group more diffusely coupled, strongly with the bilateral MCC, the right insula and the middle temporal (p< 0.05 FWE corrected; Table 3). Then, we explored the functional connectivity in both groups during the angry session and the angry grasping with the RMCC as seed region. The healthy group showed a higher coupling between the RMCC and the left SMA, (p< 0.05 FWE; Table 3). In contrast, no activation reached the statistical significance in the patients group. On the contrary, in the joyful context the right RMCC showed a functional coupling with the precentral gyrus, superior parietal lobule/SPL, posterior cingulate cortex/PCC, and the right caudate in the patients group, whereas with the left posterior insula in the healthy group.

**Table 3:**
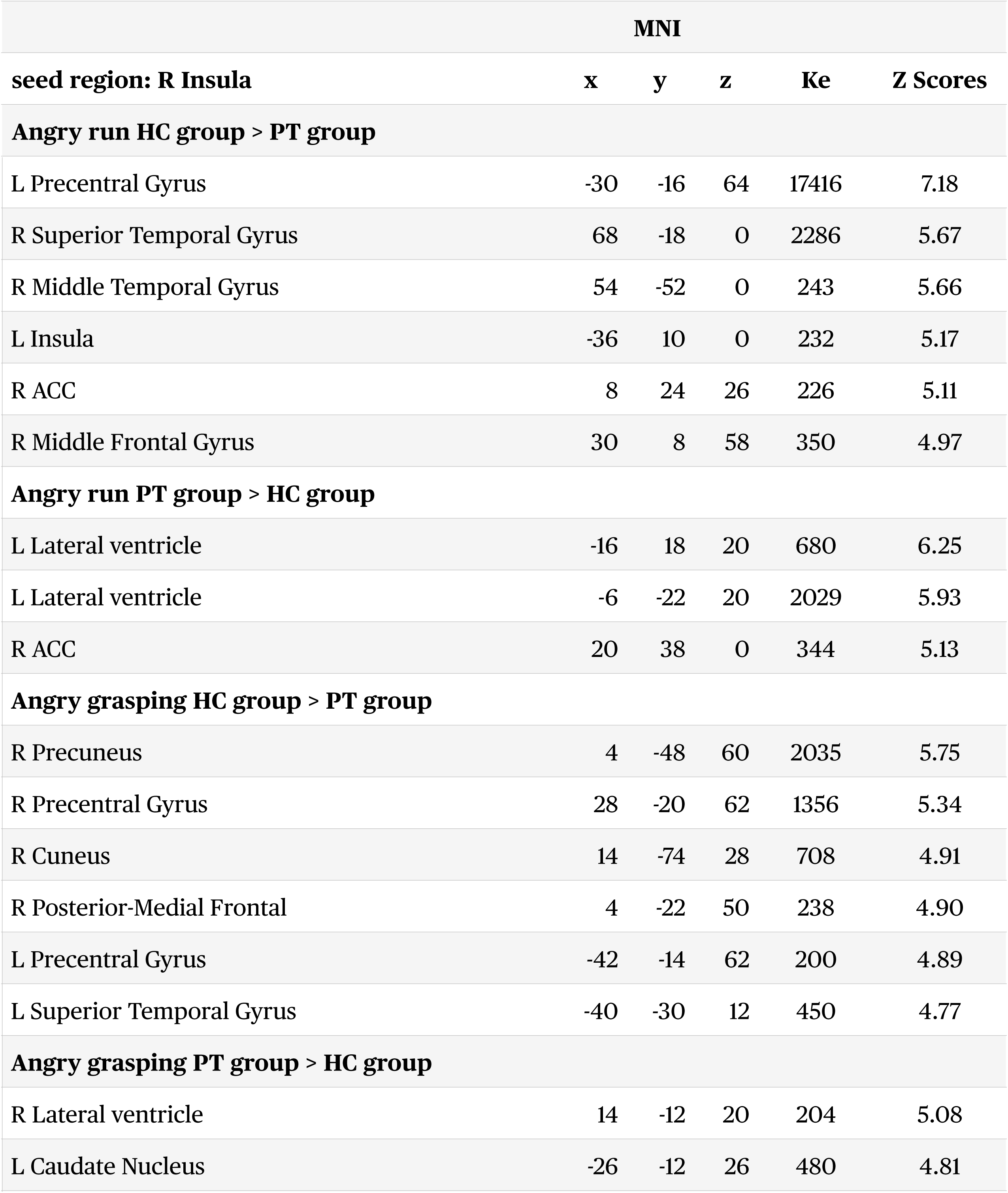
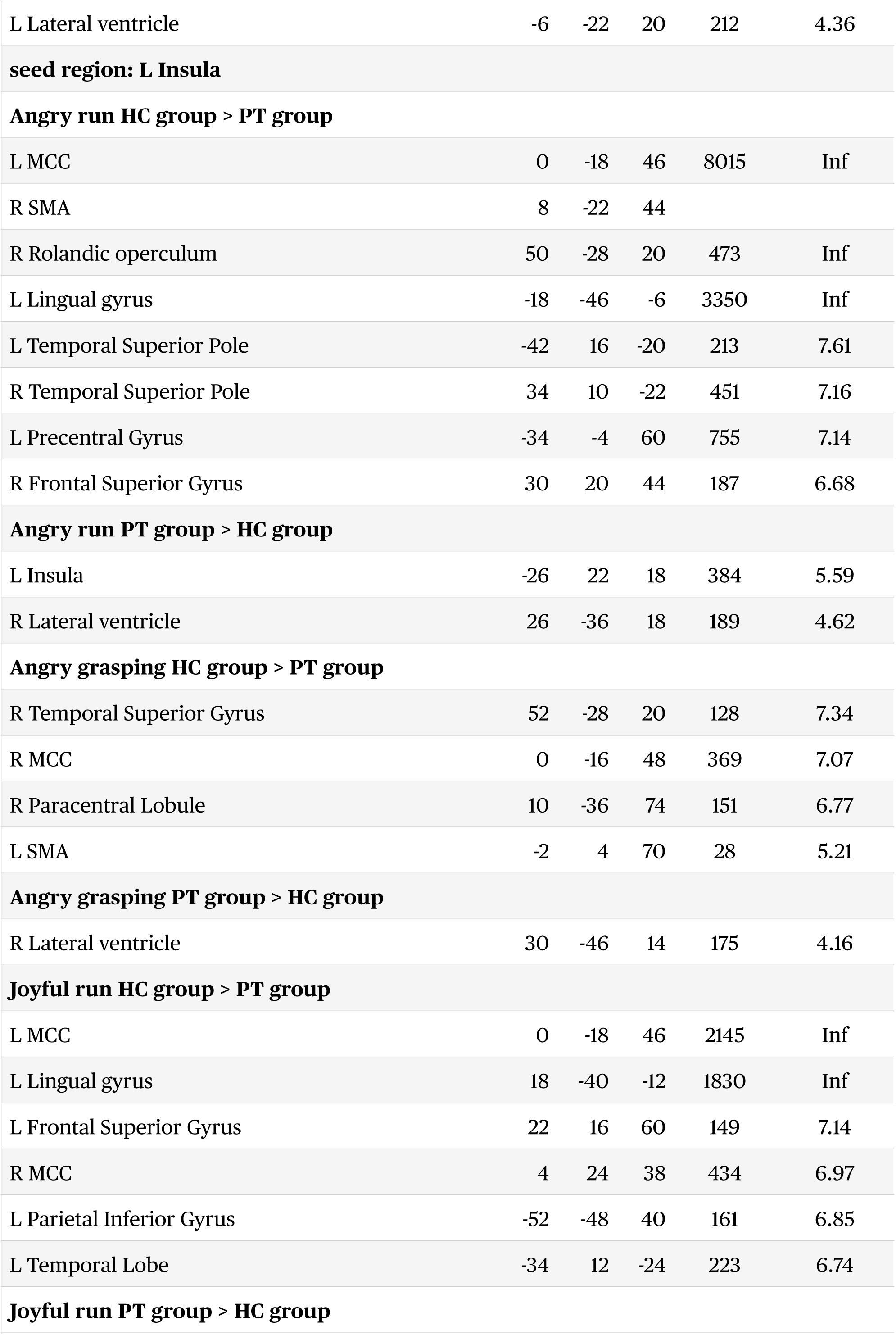
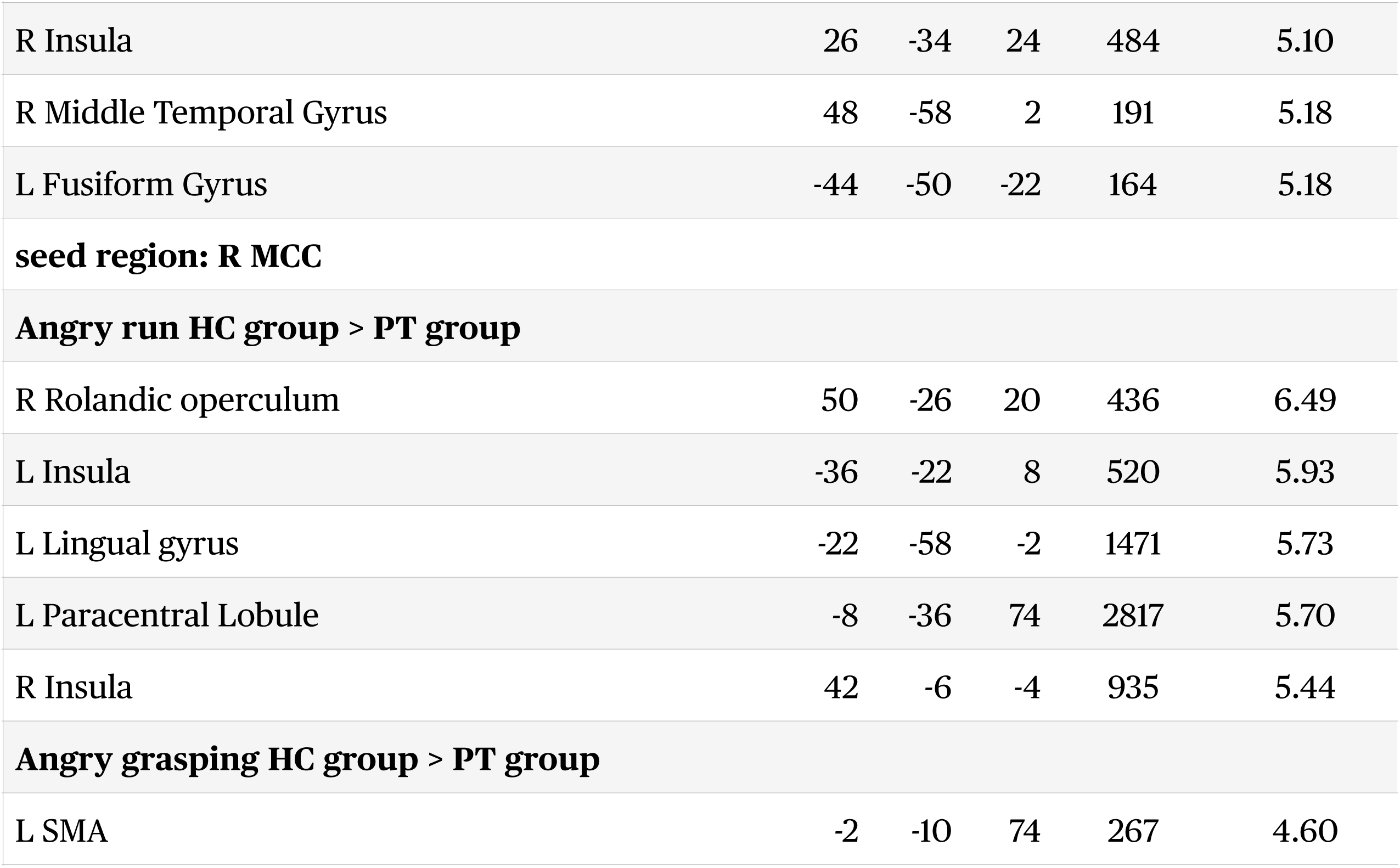
PPI Analysis. P<0.05 FWE.

|  | MNI |  |  |  |  |
| --- | --- | --- | --- | --- | --- |
| seed region: R Insula | x | y | z | Ke | Z Scores |
| **Angry run HC group > PT group** |  |  |  |  |  |
| L Precentral Gyrus | -30 | -16 | 64 | 17416 | 7.18 |
| R Superior Temporal Gyrus | 68 | -18 | 0 | 2286 | 5.67 |
| R Middle Temporal Gyrus | 54 | -52 | 0 | 243 | 5.66 |
| L Insula | -36 | 10 | 0 | 232 | 5.17 |
| R ACC | 8 | 24 | 26 | 226 | 5.11 |
| R Middle Frontal Gyrus | 30 | 8 | 58 | 350 | 4.97 |
| **Angry run PT group > HC group** |  |  |  |  |  |
| L Lateral ventricle | -16 | 18 | 20 | 680 | 6.25 |
| L Lateral ventricle | -6 | -22 | 20 | 2029 | 5.93 |
| R ACC | 20 | 38 | 0 | 344 | 5.13 |
| **Angry grasping HC group > PT group** |  |  |  |  |  |
| R Precuneus | 4 | -48 | 60 | 2035 | 5.75 |
| R Precentral Gyrus | 28 | -20 | 62 | 1356 | 5.34 |
| R Cuneus | 14 | -74 | 28 | 708 | 4.91 |
| R Posterior-Medial Frontal | 4 | -22 | 50 | 238 | 4.90 |
| L Precentral Gyrus | -42 | -14 | 62 | 200 | 4.89 |
| L Superior Temporal Gyrus | -40 | -30 | 12 | 450 | 4.77 |
| **Angry grasping PT group > HC group** |  |  |  |  |  |
| R Lateral ventricle | 14 | -12 | 20 | 204 | 5.08 |
| L Caudate Nucleus | -26 | -12 | 26 | 480 | 4.81 |
| L Lateral ventricle | -6 | -22 | 20 | 212 | 4.36 |
| **seed region: L Insula** |  |  |  |  |  |
| **Angry run HC group > PT group** |  |  |  |  |  |
| L MCC | 0 | -18 | 46 | 8015 | Inf |
| R SMA | 8 | -22 | 44 |  |  |
| R Rolandic operculum | 50 | -28 | 20 | 473 | Inf |
| L Lingual gyrus | -18 | -46 | -6 | 3350 | Inf |
| L Temporal Superior Pole | -42 | 16 | -20 | 213 | 7.61 |
| R Temporal Superior Pole | 34 | 10 | -22 | 451 | 7.16 |
| L Precentral Gyrus | -34 | -4 | 60 | 755 | 7.14 |
| R Frontal Superior Gyrus | 30 | 20 | 44 | 187 | 6.68 |
| **Angry run PT group > HC group** |  |  |  |  |  |
| L Insula | -26 | 22 | 18 | 384 | 5.59 |
| R Lateral ventricle | 26 | -36 | 18 | 189 | 4.62 |
| **Angry grasping HC group > PT group** |  |  |  |  |  |
| R Temporal Superior Gyrus | 52 | -28 | 20 | 128 | 7.34 |
| R MCC | 0 | -16 | 48 | 369 | 7.07 |
| R Paracentral Lobule | 10 | -36 | 74 | 151 | 6.77 |
| L SMA | -2 | 4 | 70 | 28 | 5.21 |
| **Angry grasping PT group > HC group** |  |  |  |  |  |
| R Lateral ventricle | 30 | -46 | 14 | 175 | 4.16 |
| **Joyful run HC group > PT group** |  |  |  |  |  |
| L MCC | 0 | -18 | 46 | 2145 | Inf |
| L Lingual gyrus | 18 | -40 | -12 | 1830 | Inf |
| L Frontal Superior Gyrus | 22 | 16 | 60 | 149 | 7.14 |
| R MCC | 4 | 24 | 38 | 434 | 6.97 |
| L Parietal Inferior Gyrus | -52 | -48 | 40 | 161 | 6.85 |
| L Temporal Lobe | -34 | 12 | -24 | 223 | 6.74 |
| **Joyful run PT group > HC group** |  |  |  |  |  |
| R Insula | 26 | -34 | 24 | 484 | 5.10 |
| R Middle Temporal Gyrus | 48 | -58 | 2 | 191 | 5.18 |
| L Fusiform Gyrus | -44 | -50 | -22 | 164 | 5.18 |
| **seed region: R MCC** |  |  |  |  |  |
| **Angry run HC group > PT group** |  |  |  |  |  |
| R Rolandic operculum | 50 | -26 | 20 | 436 | 6.49 |
| L Insula | -36 | -22 | 8 | 520 | 5.93 |
| L Lingual gyrus | -22 | -58 | -2 | 1471 | 5.73 |
| L Paracentral Lobule | -8 | -36 | 74 | 2817 | 5.70 |
| R Insula | 42 | -6 | -4 | 935 | 5.44 |
| **Angry grasping HC group > PT group** |  |  |  |  |  |
| L SMA | -2 | -10 | 74 | 267 | 4.60 |

The handgrip task results showed a significant difference between groups. During the “on” condition the patients group activated more the right insula compared to the healthy group. In contrast, no activation reached the statistical significance in the “off” contrast. On the other hand the healthy group activated more the left medial frontal gyrus, the right anterior insula, the right putamen and the right caudate in the “off” condition compared to the patients group (p< 0.05 FWE corrected; Table 4). Interesting to note, the patients group activated a more extended motor network in the “on” condition, whereas the healthy participants only the precentral gyrus. Moreover, the analyses of the PPG during the handgrip task of the patients group showed a sympathetic function generally higher than healthy control through a higher BPM mean (t=3.54 p<0.001), and a higher minimum level of isometric pressure on the device (t=2.19 p<0.035), irrespective of gender and age.

**Table 4:** Handgripping task. Whole-brain General Linear Model analyses. P<0.05 FWE.

|  | MNI |  |  | Ke | Z Scores |
| --- | --- | --- | --- | --- | --- |
|  | x | y | z |  |  |
| On PT group > HC group |  |  |  |  |  |
| R Middle Temporal Gyrus | 63 | -7 | -13 | 184 | 4.64 |
| R anterior Insula | 45 | 14 | -16 |  | 4.46 |
| L Lingual Gyrus | -30 | -82 | -10 | 75 | 4.12 |
| Off HC group > PT group |  |  |  |  |  |
| L Middle Temporal Gyrus | -63 | -34 | -4 | 5508 | 6.10 |
| R Inferior Frontal Gyrus | 39 | 26 | 2 | 155 | 5.31 |
| HC group ON > OFF |  |  |  |  |  |
| L Precentral Gyrus | -33 | -22 | 56 | 183 | 5.00 |
| PT group ON > OFF |  |  |  |  |  |
| L Posterior Central Gyrus | -42 | -28 | 56 | 930 | 6.72 |
| L SMA | 6 | -7 | 56 |  | 4.20 |
| R ACC | 12 | 50 | 32 | 864 | 5.20 |
| L ACC | -15 | 50 | 14 |  |  |
| R cerebellum 4_5 | 15 | -52 | -19 | 195 | 5.07 |
| L Frontal Superior Gyrus | -30 | 23 | 47 | 75 | 3.97 |

The BOLD resting state analysis did not show significant differences between groups in connectivity in the default mode, salience, and insular cortex networks, as well as ASL resting state in the insula.

## Discussion

We studied the role of the insular cortex in cardiovascular control using a multimodal method consisting of emotional and motor tasks, as well as resting state tasks during fMRI session, in patients with different types of heart arrhythmias compared to healthy controls.

The present findings confirm our hypothesis. First, what is evident in the healthy controls is the key involvement of the RMCC and its strongly functional coupling mostly with the SMA in anger processing. In line with our previous findings, this indicates an emotional processing facing angry situations that prompts essentially to act (Mazzola et al. 2016, 2020). On the other hand, the greater generalized involvement of the bilateral posterior insulae in the patients group reveals an emotional processing in the same situations that instead prompts essentially an automatic engagement of the sympathetic nervous system/SNS and, accordingly, of the bodily reactions. Specifically, in facing someone grasping simple objects with an angry expression, they showed the involvement of the left insula, previously associated with the cardiac-related pulsatility (Atwi et al. 2020). Moreover, beyond our expectations, the bilateral insulae were consistently functionally coupled with the bilateral lateral ventricles under both the angry grasping and the angry context, but not under the joyful context. These coupled dynamics suggest that insular activity may act as an additional driver of macroscopic variation of cerebrospinal fluid/CSF flow due to the neurovascular coupling. In fact, recent reports suggest a possible link between CSF pulsations and cerebral neural activity (Fultz et al., 2019; van Veluw et al., 2020), as well as sympathetic tone (Attarpour et al., 2021; Picchioni et al., 2022). In addition, a recent study has explored the relationship between atrial fibrillation/AF, glymphatic dysfunction, and CSF dynamics (Hofer et al. 2025). Even the motor task performed by the handgrip exercise exhibited results that go in the same direction. Indeed, they revealed an overall increased sympathetic activation both at the brain and peripheral levels in the patients during the physical effort while squeezing the hand device.

All these brain noteworthy results about the insular activity and connectivity cannot be associated with or could be better explained by an a priori different baseline neural activity of the insula, its functional connectivity and blood flow. The BOLD and ASL resting state results did not find any statistically significant differences between the two groups. Instead, the fact that the insula activation in the patients groups was statistically significant at a very lower threshold after the physiological noise correction of fMRI data using peripheral measures of respiration and cardiac pulsation, points out the functional, i.e. contextual, nature of this emotional-physiological interaction in our patients. In addition, the blood test results showed no significant differences between groups in catecholamines level, supporting at least in our groups the contextual quality of the reactivity of the insula and sympathetic tone, therefore not otherwise explained by this factor. On the other hand, the significant differences in others cytokines levels may be related with the inflammatory background of these patients as often due to the CVD diagnosis as an effect of elevated reactivity of the SNS (Lazzerini et al. 2023). Unfortunately, our blood test samples were limited due to technical problems.

Taken together, these brain findings unravel an anger-insula-heart link that can be automatically triggered by the experiencing of an angry vs joyful situation in our participants with different types of heart arrhythmias.

From a psychological point of view, all our participants did not present clinical symptoms, neither an a priori angry-aggressive profile, personality traits or exceptional motivations grounding their anger or mild irritation in personal life. Nevertheless, some clear differences stand out in anger management between the two groups. Indeed, compared to the healthy participants, the patients struggle and attempt more to curb and monitor themselves to prevent any explosive manifestations of their anger toward the others, evidently feeling the emotional bodily pressure increased and increasing. In doing so, on one hand, we may better realize how much that expose them to a sympathetic hyperactivity enhancing the risk of having or worsening the CVD symptoms, but also to the whole physiological implications correlated, for instance, the cytokines storms and the neuro-inflammatory responses. From the other hand, it exposes them to live non-traumatic life events as more “stressful”, i.e. with an intense emotional-sympathetic pressure, centralizing the link between emotional management and the chronicity of symptoms or diseases.

To our knowledge, this is the first human brain imaging study to investigate this anger-insula link in persons with CVDs with a multimodal approach. It helps to unveil how the cardiovascular brain circuit is engaged by anger processing at least in these different types of heart arrhythmias. Nevertheless, there are some limitations. Clearly, the homogeneity of diagnosis of the clinical group may not be sufficient, and we tried to control the variability of the sample at least by matching of age and gender. Moreover, the limited number of blood samples did not allow a more in-depth analysis of sympathetic activation in terms of inflammatory markers. Further investigations are required.

In conclusion, since the CVDs are not necessarily explained by the psychopathology, but maybe more by the personal story, and nor being anyone of us immune from them, understanding this anger-insula-heart link may factually contribute to the prevention, i.e. lower the probability, of adverse cardiac outcomes, as well as to the necessary treatment of the person, taking into account the emotion-related cardiovascular pathologies and the psychological health (Levine et al. 2021; Arciero et al. 2018).

## Acknowledgements

Pr Haran Burri and Pr François Mach, Cardiology service of the University Hospitals of Geneva, Pr Nicolas Vuilleumier and Dr Sabrina Pagano for the blood testing, Bruno Bonnet the Lab manager of the Brain and Behavior Laboratory (BBL) of the University of Geneva for the technical support of the fMRI. All the data have been collected from March 2017 to December 2019 at the Liaison Psychiatry service of the University Hospitals of Geneva.

## Funding Statement

This research received no specific grant from any funding agency, commercial or not-for-profit sectors. The funding of the study was provided by Pr Guido Bondolfi’s university account and the Liaison Psychiatry service of the University Hospitals of Geneva.

## Data Availability Statements

The data underlying this article were provided through the University Hospitals of Geneva. Data will be shared on reasonable request to the corresponding author with permission of the University Hospitals of Geneva.

## Notes

### Competing Interest Statement

The authors have declared no competing interest.

